# Preferential lipolysis of DGAT1 over DGAT2 generated triacylglycerol in Huh7 hepatocytes

**DOI:** 10.1101/2021.06.18.449045

**Authors:** Rajakumar Selvaraj, Sarah V. Zehnder, Russell Watts, Jihong Lian, Randal Nelson, Richard Lehner

**Author notes:** <u>Correspondence:</u>.

## Abstract

Hepatic steatosis is defined by accumulation of neutral lipids in lipid droplets (LDs) including triacylglycerol (TG) and steryl esters. Two distinct diacylglycerol acyltransferases (DGAT1 and DGAT2) catalyze synthesis of TG in hepatocytes. TG formed through either DGAT1 or DGAT2 appears to be preferentially directed to distinct intercellular fates, such as fatty acid production for oxidation or very-low density lipoprotein assembly, respectively. Because of the preferential use of TG generated by DGAT1 and DGAT2, we hypothesized that targeting/association of lipolytic machinery to LDs would differ depending on whether the TG stores were generated through DGAT1 or DGAT2 activities. Inhibition of DGAT1 or DGAT2 in human hepatoma cells (Huh7) incubated with oleic acid resulted in only a small change in TG accretion suggesting that the two DGATs can compensate for each other in fatty acid esterification. This compensation was not accompanied by changes in *DGAT1* or *DGAT2* mRNA expression. DGAT1 inhibition (TG synthesized by DGAT2) resulted in large LDs, whereas DGAT2 inhibition (TG synthesized by DGAT1) caused the accumulation of numerous small LDs. Oleic acid treatment increased mRNA and protein expression of the LD-associated protein PLIN2 but not PLIN5 or the lipase ATGL and its activator ABHD5/CGI-58. Inactivation of DGAT1 or DGAT2 did not alter expression (mRNA or protein) of ATGL, ABHD5/CGI-58, PLIN2 or PLIN5, but inactivation of both DGATs increased PLIN2 abundance despite a dramatic reduction in the number of LDs. ATGL localized preferentially to DGAT1-made LDs rather than to DGAT2-made LDs, and TG in these LDs was preferentially used for fatty acid (FA) oxidation. A combination of DGAT2 inhibitor and the pan lipase inhibitor E600 resulted in large LDs, suggesting that the small size of DGAT1-made LDs is due to a lipolytic process.

## Introduction

Neutral lipids such as triacylglycerol (TG) and steryl esters are stored in ubiquitous storage organelles called lipid droplets (LDs) (Walther et al., 2017). Lipids in LDs serve as reservoirs of energy, as well as provide substrates for membrane synthesis and ligands for cellular signaling (Walther and Farese, 2012; Wilfling et al., 2014). The process of deposition of lipids in LDs protects cells from deleterious effects of excessive accumulation of lipotoxic fatty acids (Koliwad et al., 2010; Listenberger et al., 2003). The buildup of LDs in various pathological states, including nonalcoholic fatty liver disease (NAFLD) is well documented (Seebacher et al., 2020) but the homeostatic mechanisms governing LD formation and turnover have not yet been fully elucidated.

Acyl-CoA: diacylglycerol acyltransferases (DGAT) 1 and 2 catalyze TG synthesis through FA esterification to diacylglycerol (Cases et al., 1998; Yen et al., 2008). The two DGATs belong to separate protein families, share no sequence homology, and have distinct physiological functions, as demonstrated by different phenotypes of *Dgat1*^-/-^ and *Dgat2*^-/-^ mice (Smith et al., 2000; Stone et al., 2004). Both DGAT1 and DGAT2 are localized in the endoplasmic reticulum (ER), while DGAT2 may also interact with the LD surface (Stone et al., 2009). DGAT1 and DGAT2 have been reported to utilize both exogenously supplied or endogenously synthesized fatty acids (Li et al., 2015; Villanueva et al., 2009; Wurie et al., 2012).

Mice lacking DGAT1 are resistant to diet-induced obesity, while mice overexpressing DGAT1 mice become more obese than wild type mice when fed high-fat diets (Chen and Farese, 2005; Chen et al., 2003; Chen et al., 2002). DGAT2 can partially compensate for the loss of DGAT1, while DGAT1 does not appear to have the ability to compensate for the loss of DGAT2 because global DGAT2 knockout mice have drastically reduced TG content and die shortly after birth (Stone et al., 2004). DGAT2 was proposed to be localized at the bridge between the ER and LD and therefore is exquisitely positioned to promote LD expansion (McFie et al., 2018). Inactivation of DGAT2 in primary mouse hepatocytes restricts LD expansion, leading to generation of small-sized LDs via DGAT1 activity (Li et al., 2015). Various acyltransferases, fatty acid activating enzymes, lipases and regulatory proteins interact with the LD surface (Walther and Farese, 2009), which leads to remodeling of the LD lipid content through a cycle of neutral lipid turnover and (re)synthesis (Paar et al., 2012; Schott et al., 2019). Perilipins (PLINs) are well-studied LD surface proteins which serve unique structural and regulatory functions in LDs, including regulating access and activity of lipases. In hepatocytes, one of the well-studied lipases that hydrolyzes LD-associated TG is adipose triglyceride lipase (ATGL). PLIN1, PLIN2, and PLIN5 dampen ATGL activity by either reducing access of ATGL to LDs or by sequestering the ATGL activator ABHD5/CGI-58 (Granneman et al., 2009; Wang et al., 2011; Zimmermann et al., 2004). In this study we tested the hypothesis that targeting/association of the lipolytic machinery to LDs would differ depending on whether the TG stores were generated through DGAT1 or DGAT2 activities.

## Materials and methods

### Cell culture and reagents

Huh7 (human hepatocellular carcinoma) cells were cultured at 37°C and 5% CO_2_ in Dulbecco’s Modified Eagle’s Medium (DMEM) containing 10% fetal bovine serum, penicillin/streptomycin and supplemented with 1.0 mM pyruvate, 2.0 mM L-glutamine and 25 mM glucose (Invitrogen Canada, Burlington, ON, Canada). The medium was refreshed daily. Cells were passaged once they reached 80% confluence by digestion with a 0.25% trypsin EDTA solution. Fatty acid-free bovine serum albumin (BSA) and Complete Protease Inhibitor cocktail tablets were procured from Roche Diagnostics (Laval, QC, Canada). BODIPY 493/503 and LipidTox Red were purchased from Invitrogen (Carlsbad, CA). [9,10(n)-^3^H]oleic acid (OA) (54.6 Ci/mmol) and [^14^C]glycerol (250 μCi/mmol) were from Perkin Elmer. Inhibitors of DGAT1 (PF-04620110), and DGAT2 (PF-06424439) were synthesized by Pfizer Inc. (Groton, CT) and obtained from Sigma-Aldrich. Diethyl-p-nitrophenyl phosphate (E600) was obtained from Sigma-Aldrich. All other reagents were of analytical grade or higher.

### Determination of cellular TG

Gas chromatography was used to quantify the cellular mass of TG. The analysis was carried out on the Agilent Technologies 6890 Series gas chromatograph equipped with a flame ionization detector (Palo Alto, CA). Samples were injected onto an Agilent high performance capillary column (HP-5, 15 m × 0.32 mm × 0.25 μm). The oven temperature was raised from 170 to 290°C at 20°C/min and then to 340°C at 10°C/min, with helium as a carrier gas and a constant flow rate of 4.5 ml/min. In some experiments, TG content in cells and media was determined using “Infinity” TG reagent (Sigma Diagnostics, Inc.) using authentic TG (triolein) as the internal standard.

### Detection of LDs by confocal fluorescence scanning microscopy

Huh7 cells were preincubated for 1 h with/without 5 μM DGAT1 and/or DGAT2 inhibitors in serum-free medium, followed by 4 h incubation in DMEM containing 0.4 mM OA/0.5% BSA ± DGAT inhibitors. Following the incubation, the cells were washed three times with phosphate-buffered saline (PBS), fixed with 4% formaldehyde for 20 minutes, followed by three washes with PBS. LDs were labeled with 2 μg/mL BODIPY 493/503 in PBS for 30 minutes. Samples were mounted using Vectashield mounting medium for fluorescence with DAPI (Vector Laboratories Inc. Cat# H-1200). Images were captured by a spinning disk confocal microscope (Olympus IX81-DSU). The quantification of LD numbers and sizes were determined using Fiji-ImageJ.

### Immunoblotting

Huh7 cells were harvested in ice-cold IM buffer (250 mM sucrose, 50 mM Tris, 1 mM EDTA, pH7.4), sonicated and protein concentration was determined. Proteins were separated by SDS-PAGE, transferred to Immobilon-P membrane (PVDF) (catalog #IPVH00010; Millipore) and subjected to immunoblotting. The following primary antibodies (diluted in 3% w/v BSA in TBST) were used: adipose triglyceride lipase (ATGL) (1:1000, Cell Signaling #2138), perilipin5 (PLIN5) (1:2000, PROGEN #GP31), acyl-CoA synthetase long-chain family member 1 (ACSL1) (1:1000, Cell Signaling, #4047), perilipin 2 (PLIN2) (1:1000, Abcam #108323), calnexin (Cnx) (1:1000, Enzo Life Sciences #SPA-865), glyceraldehyde 3-phosphate dehydrogenase (GAPDH) (1:5000, Abcam #ab8245). Membranes were incubated with primary antibodies overnight (12-16 h) at 4°C. After washing, the membranes were incubated with appropriate secondary antibodies, diluted to 1:5000 in 5% w/v skimmed milk in TBST, for 1 h at room temperature. The secondary antibodies were HRP-labelled goat anti-rabbit IgG (Invitrogen, #31460) or HRP-labelled goat anti-mouse IgG (Invitrogen, #31460). Immunoreactive proteins were detected by enhanced chemiluminescence (Classico substrate, Immobilon-Millipore, USA, #WBCUC0500) and visualized by G: BOX system (SynGene, UK). The relative intensities of immunoreactivity were analyzed by densitometry using the GeneTools program (SynGene).

### Double-label pulse-chase experiments to assess cellular TG synthesis and turnover

Huh7 cells were preincubated ± DGAT inhibitors for one hour with serum free DMEM media. After preincubation the cells were pulse labelled for 4 h in 2 ml of DMEM containing DGAT1 or DGAT2 inhibitors, 5 μCi [^3^H]OA (Perkin Elmer) + 0.4 mM OA/0.5% BSA complex, 1 μCi [^14^C]glycerol (Perkin Elmer) + 50 μM glycerol to stimulate glycerolipid synthesis. Some of the dishes were harvested at the end of the 4 h pulse. Remaining cells were washed three times with DMEM containing 0.5% BSA, changed to chase DMEM serum free medium with 5 μM DGAT1&2 inhibitors to prevent re-esterification with or without 200 μM E600 (pan-lipase inhibitor), and incubated for 6 h. Cells were harvested in PBS, lipids were extracted by modified Folch method (Folch et al., 1957) and subsequently separated by thin layer chromatography (TLC) with the solvent system heptane:isopropyl ether:acetic acid (15:10:1). Lipids were visualized by exposure to iodine, recovered from the TLC plates and the radioactivity was determined by liquid scintillation counting. The percentage of lipid turnover was expressed relative to radioactivity obtained in the pulse. Chase media obtained from the metabolic labelling studies were analyzed for acid-soluble metabolites (ASM) resulting from fatty acid β-oxidation. Thirty μl of 20% BSA and 16 μl of 70% perchloric acid were added to 200 μl of culture medium from each sample. Media were then centrifuged at 25,000g for 5 min before an aliquot of the supernatant was counted for radioactivity.

### Localization of LD-associated proteins

#### Generation of ATGL-EGFP, ATGL(S47A)-EGFP, PLIN2-EGFP, ABHD5/CGI-58-mCherry and PLIN5-mCherry

ATGL-GFP was created by inserting mouse ATGL cDNA (a generous gift from Rudolf Zechner, University of Graz) into the AGE I and Bgl II sites of pEGFP-C1. The catalytically inactive ATGL(S47A) mutant was was made by site directed mutagenesis using the QuickChange II kit (Stratagene). PLIN2-GFP was a generous gift from Carole Sztalryd (University of Maryland). PLIN5-mCherry was constructed by replacing the EYFP sequence of pEYFP-C1-PLIN5 (a generous gift from Carole Sztalryd, University of Maryland) with mCherry. ABHD5/CGI-58-mCherry was constructed by inserting mouse ABHD5/CGI-58 cDNA (a generous gift from Dawn Brasaemle, Rutgers University) downstream of mCherry.

#### Visualization of intracellular localization of LD proteins

Huh7 cells were plated in 12-well plates with coverslips in complete medium (DMEM with 10% FBS plus penicillin and streptomycin). Cells at 40-60% confluency were then transfected with various LD protein constructs including full length ATGL-EGFP, inactive point mutated ATGL(S47A)-EGFP, ABHD5/CGI58-mCherry, PLIN2-EGFP and PLIN5-mCherry using Lipofectamine 2000 transfection reagent (Thermo Fisher Scientific) according to the manufacturer’s instructions. After 24 h incubation, the medium was changed to preincubation of serum free DMEM medium with/without 5 μM DGAT1 or DGAT2 inhibitors for 1 h. After preincubation, medium was replaced with serum free DMEM with/without DGAT1 or DGAT2 inhibitors plus 0.4 mM OA/0.5% BSA, and cells were incubated for 4 h. Cells were then washed with PBS (3×5min), fixed with 4% formaldehyde, and then stained for 30 min with LipidTox Red (1:1000 dilution) or for 15 min with BODIPY 493/503 (2 μg/ml) (Invitrogen) to visualize LDs. Mounting medium with DAPI (VECTASHIELD, Vector Laboratories) was used to preserve specimens. Images were acquired using an Olympus IX-81 CSU10 spinning disk confocal microscope.

### RNA isolation and real-time qPCR analysis

Huh7 cells were pre-incubated for 1 h in serum-free DMEM medium with/without DGAT1 or DGAT2 inhibitors, followed by a 4 h incubation in DMEM + 0.4 mM OA complexed with 0.5% BSA in the presence/absence of the inhibitors. Total RNA was then isolated using TRIZOL Reagent according to the manufacturer’s instructions. First-strand cDNA was synthesized from 2 μg of total RNA using Superscript III reverse transcriptase (Invitrogen) primed by oligo (dt)12-18 (Invitrogen) and random primers (Invitrogen). Real-time quantitative qPCR was performed with Power SYBR® Green PCR Master Mix kit (Life Technologies, UK) using the StepOnePlus real-time PCR system (Applied Biosystems, Canada). All primers were synthesized by Integrated DNA Technologies (Canada). Primers for genes tested are listed in Supplemental Table S1. Data were analyzed with the StepOne software (Applied Biosystems). Standard curves were used to calculate mRNA abundance relative to that of the control gene *PPIA* encoding cyclophilin A.

### Statistical analysis

Data are plotted as mean ± SEM. Significant differences between two groups were determined by unpaired two-tailed t-tests. Data from studies using Huh7 cells incubated with DGAT1 or DGAT2 inhibitors were analyzed by two-way ANOVA followed by Bonferroni post hoc tests (Graph Pad PRISM 8 software). Differences were considered statistically significant at *P < 0.05, **P < 0.01, ***P < 0.001 and #P<0.0001.

## Results

### DGAT1 and DGAT2 can compensate for each other to synthesize TG but give rise to LDs with different morphology

We determined whether or not the inhibition of individual DGAT enzymes affects TG synthesis and LD formation in human hepatocytes (Huh7 cells). We confirmed both *DGAT1* and *DGAT2* mRNA are expressed in Huh7 cells (Figure 1A). Inhibition of individual or both DGATs together did not result in any compensatory increase of expression of either *DGAT* gene (Figure 1A). We then investigated the potential impact of acute inhibition of DGAT1 or DGAT2 on esterification and storage of exogenously supplied OA in Huh7 cells. As expected, OA supplementation significantly increased TG concentration in Huh7 cells (Figure 1B). Treatment of Huh7 cells with individual inhibitors of DGAT1 or DGAT2 did not significantly reduce OA incorporation into cellular TG, but the presence of both DGAT1 and DGAT2 inhibitors resulted in prevention of TG accretion (Figure 1B), indicating the potency of both DGAT1 and DGAT2 inhibitors and compensation of one DGAT enzyme for the loss of the other in the esterification of exogenously supplied fatty acid. We further analyzed whether inhibition of DGAT activities alters LD morphology. OA supplementation increased the size and number of LDs (Figure 1C,D). Treatment with DGAT1 inhibitor resulted in larger size LDs, but the number of LDs was reduced, whereas DGAT2 inhibition led to the formation of increased number of smaller size LDs(Figure 1C,D). Cells treated with both DGAT inhibitors showed similar phenotype as cells without OA incubation, suggesting no new LD formation (Figure 1C,D), in agreement with lipid mass data (Figure 1B).

**Figure 1:**
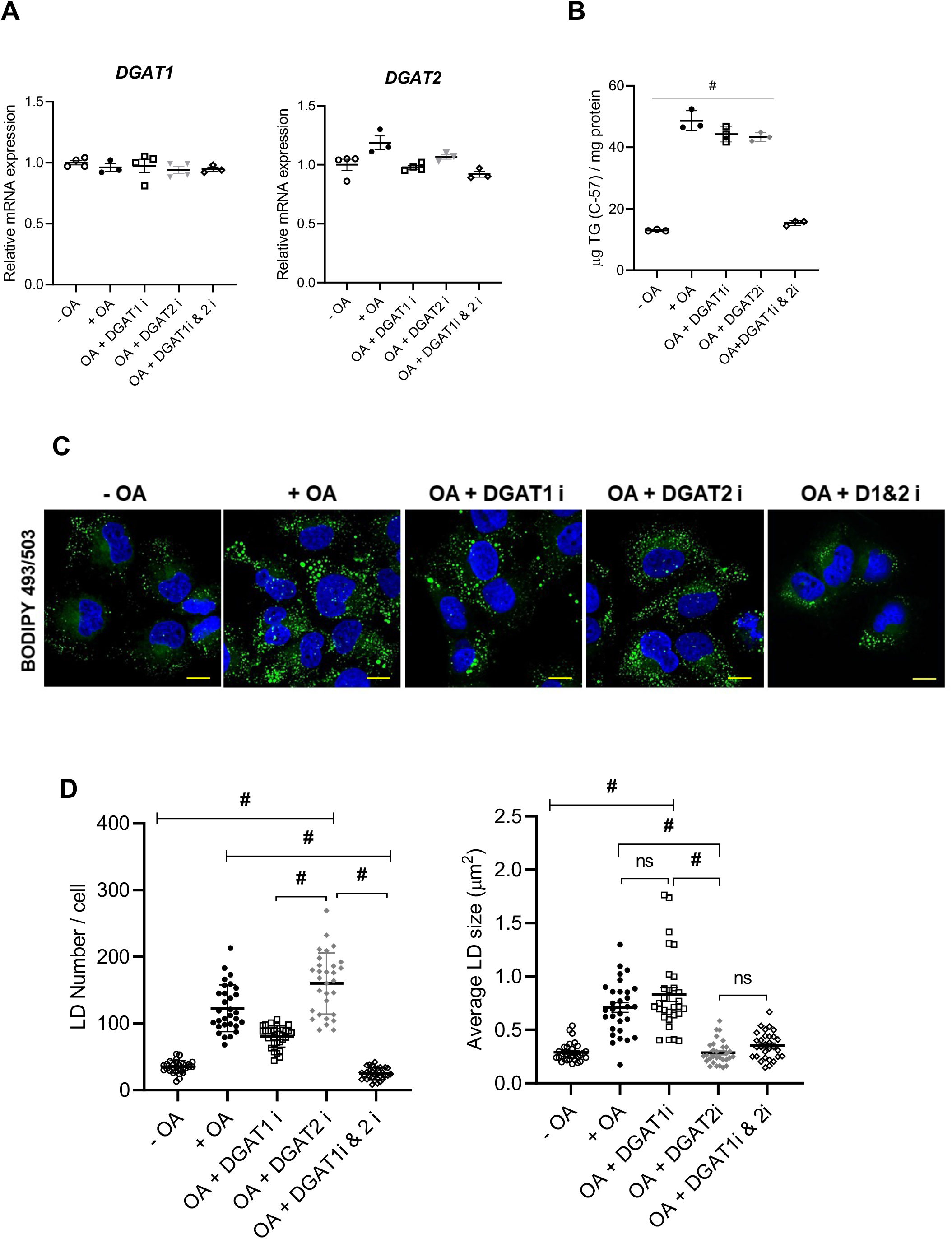
DGAT1 and DGAT2 generate different size of LDs. Huh7 hepatocytes were incubated with serum free DMEM containing DGAT1 or DGAT2 inhibitors singly or together and 0.4 mM OA/0.5% BSA complex for 4 h, followed by RNA and lipid analysis. (A) mRNA expression of *DGAT1* and *DGAT2* by qRT-PCR. (B) Intracellular TG quantified by gas chromatography. (C) LDs were visualized with BODIPY 493/503 (Green-LD, Blue-Nuclei, Scale=10 μM). (D) Quantification of LD numbers and average size of LDs represents mean ± SEM from 30 cells.

### Effect of DGAT inhibitors on lipogenic and lipolytic regulators

Next, we determined expression of specific lipid synthesis/lipolysis genes and proteins in DGAT inhibitor treated Huh7 cells. OA supplementation resulted in reduced expression of a key transcription factor regulating the expression of lipogenic genes *SREBF1* and its target *SCD* but not *FASN* (Figure 2A). Inactivation of DGAT1, DGAT2 or both DGATs did not further affect expression of *SREBF1* or *SCD* (Figure 2A). Inhibition of DGAT2 but not DGAT1 decreased the expression of *FASN* (Figure 2A). Perilipins 2 and 5 (PLIN2 and PLIN5) play an important role in the regulation of accretion and turnover of LD-associated TG in hepatocytes. Interestingly, while Huh7 cells supplemented with OA increased TG mass and expectedly PLIN2 expression (both mRNA and protein), PLIN5 abundance was unaltered even though TG synthesis and storage were increased (Figure 2B,D). Incubation with OA and treatment with DGAT inhibitors did not affect the expression of *BSCL2* encoding seipin or *FITM2* encoding fat storage-inducing transmembrane protein 2 (FIT2) (Figure S1). Expression (mRNA and protein) of *PNPLA2*, (the gene encoding ATGL, a key lipase hydrolyzing LD-associated TG), and *ABHD5* (encoding the ATGL activator ABHD5/CGI-58) were not affected by OA supplementation or DGAT inhibition (Figure 2C,D).

**Figure 2:**
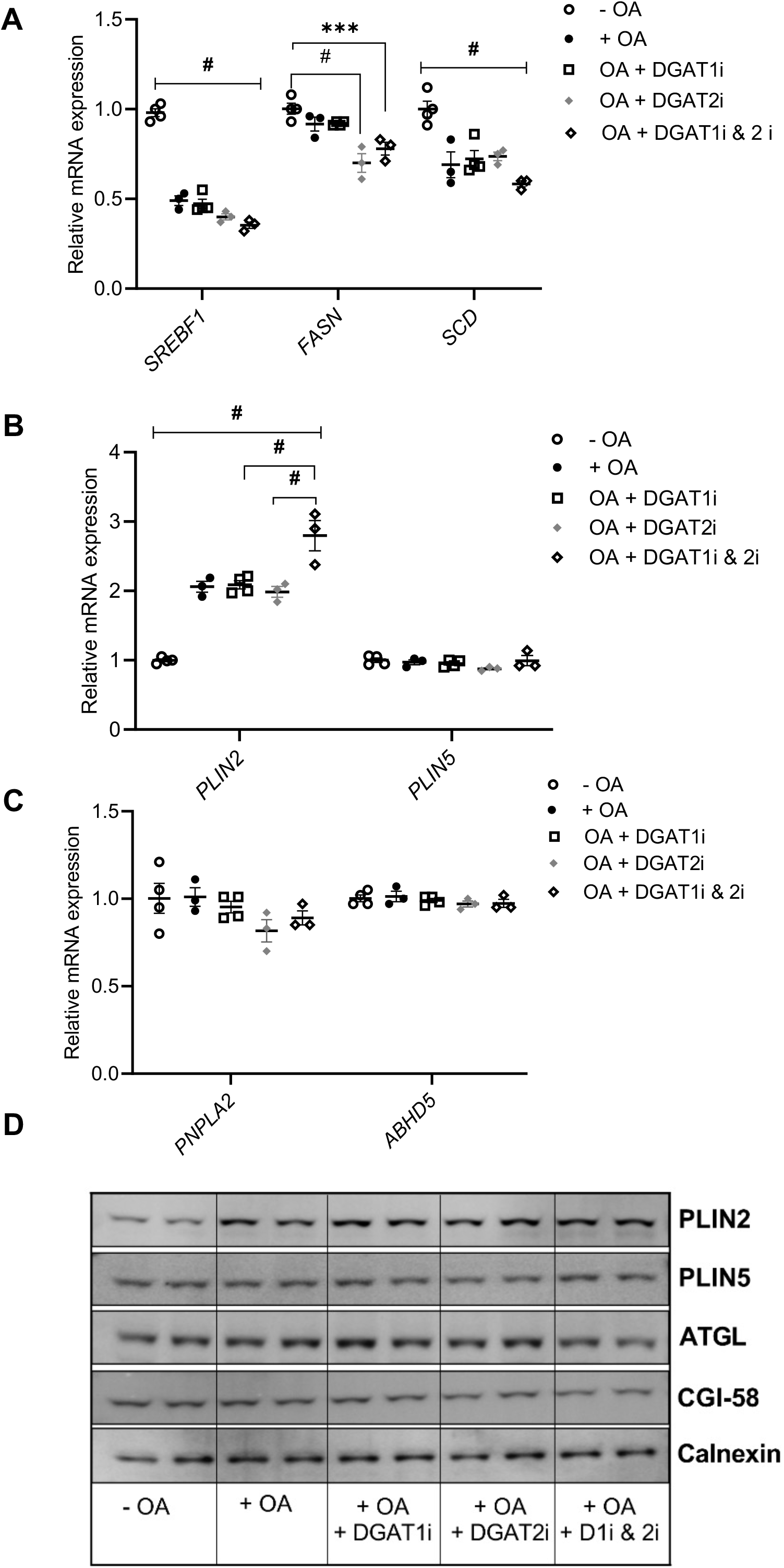
Inhibition of DGATs has different effects on expression of lipogenic and lipolytic regulators. Huh7 hepatocytes were preincubated for 1 h with serum free DMEM with DGAT1 or DGAT2 inhibitors singly or together followed by incubation with or without 0.4 mM OA/0.5% BSA complex ± DGAT inhibitors for 4 h. (**A**) mRNA expression of lipogenic genes *SREBF1, FASN* and *SCD*. (**B**) mRNA expression of *PLIN2* and *PLIN5*. (**C**) mRNA expression of *PNPLA2* and *ABHD5*. (**D**) Immunoblotting of PLIN2, PLIN5, ATGL and ABHD5/CGI-58. Calnexin served as a loading control.

### Inhibition of lipolysis affects morphology of DGAT1 generated LDs

Inhibition of DGAT2 resulted in an increased number of small sized LDs, while inactivation of DGAT1 led to a decreased number of LDs, which were larger in size. We hypothesized that small sized LDs generated by DGAT1 could arise from lipolysis. To test this hypothesis, we analyzed LD morphology in Huh7 cells treated with a pan-lipase inhibitor diethyl-*p*-nitrophenyl phosphate (E600) in the presence or absence of DGAT1 or DGAT2 inhibitors. Inhibition of lipolysis prevented increased number of small LDs generated by DGAT1 (DGAT2 inhibited) (Figure 3 A-C). No statistically significant differences in LD numbers and sizes were observed when lipolysis was inactivated in the presence of DGAT1 inhibitor (Figure 3 A-C). These results suggest that DGAT1-synthesized TG is targeted for lipolysis more effectively than DGAT2-made TG.

**Figure 3:**
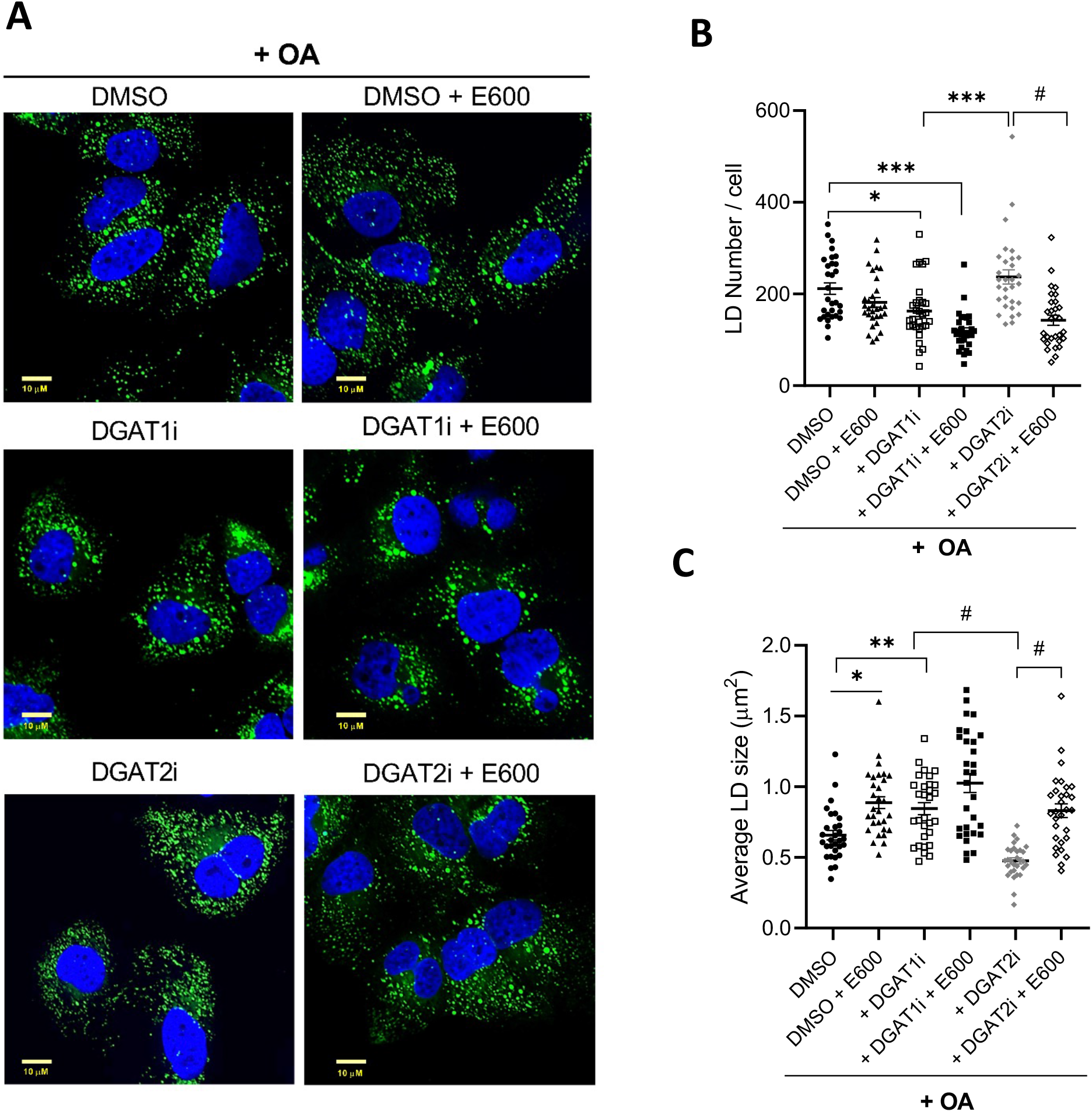
Inhibition of lipolysis influences DGAT1-generated LD size and number. Huh7 hepatocytes were preincubated for 1 h with serum free DMEM with DGAT1 or DGAT2 inhibitors singly or together followed by incubation with 0.4 mM OA/0.5% BSA complex ± DGAT inhibitors for 4 h in the presence/absence of lipase inhibitor E600. (**A**) LDs were visualized with BODIPY 493/503, nuclei with DAPI. (**B**) Quantification of LD numbers and (**C**) size of LDs, Data are mean + SEM from 30 cells.

Inactivation of lipolysis did not alter the abundance of LD-associated proteins PLIN2, ATGL, or ABHD5/CGI-58 (Figure S2) but increased LDs’ size without changing the number of LDs (Figure 3A-C). Compared to the control, the combined treatment with DGAT1 and lipase inhibitors significantly increased ATGL and ABHD5/CGI58 protein abundance (Figure S2), while LD numbers were slightly reduced (Figure 3A and B). Inactivation of lipolysis in DGAT2 inhibited cells (TG synthesis proceeding through DGAT1 catalyzed esterification) decreased the number and increased the size of LDs (Figure 3A-C) and increased PLIN2 protein abundance (Figure S2). These results suggest that the smaller size of DGAT1-made LDs results from the lipolytic process.

### Effect of DGAT inhibitors on TG turnover in hepatocytes

To determine the susceptibility of DGAT1-and DGAT2-made LDs to lipolytic turnover we employed a pulse/chase radiolabelling assay using [^14^C]glycerol to label the glycerol backbone and [^3^H]OA to label the acyl group of TG. During the subsequent 6 h chase period about 50% of the pulse label in TG (both [^3^H] and [^14^C]) was lost in control cells (Figure 4A,B) indicating a robust TG turnover rate. Incubation with a lipolysis inhibitor E600 significantly decreased TG turnover (Figure 4A,B). Inhibition of DGATs had a differential effect on TG turnover. Inhibition of DGAT1 attenuated TG turnover to a greater extent than inhibition of DGAT2, suggesting that DGAT1-made TG was turning over faster than DGAT2-made TG (Figure 4A,B). During the chase period, the only source of fatty acids for fatty acid oxidation were the fatty acids released by lipolysis of preformed TG stored in LDs because inactivation of lipolysis completely prevented fatty acid oxidation (Figure 4C). Increased turnover of DGAT1-made TG is also supported by measurement of acid soluble metabolites produced by fatty acid oxidation where more oxidation of fatty acids occurred when DGAT2 was inhibited (Figure 4C). Increased provision of DGAT1-made TG for fatty acid oxidation is further supported by elevated expression of *CPT1A*, a gene encoding a transporter of fatty acids into mitochondria, when DGAT2 was inhibited (Figure 4D).

**Figure 4:**
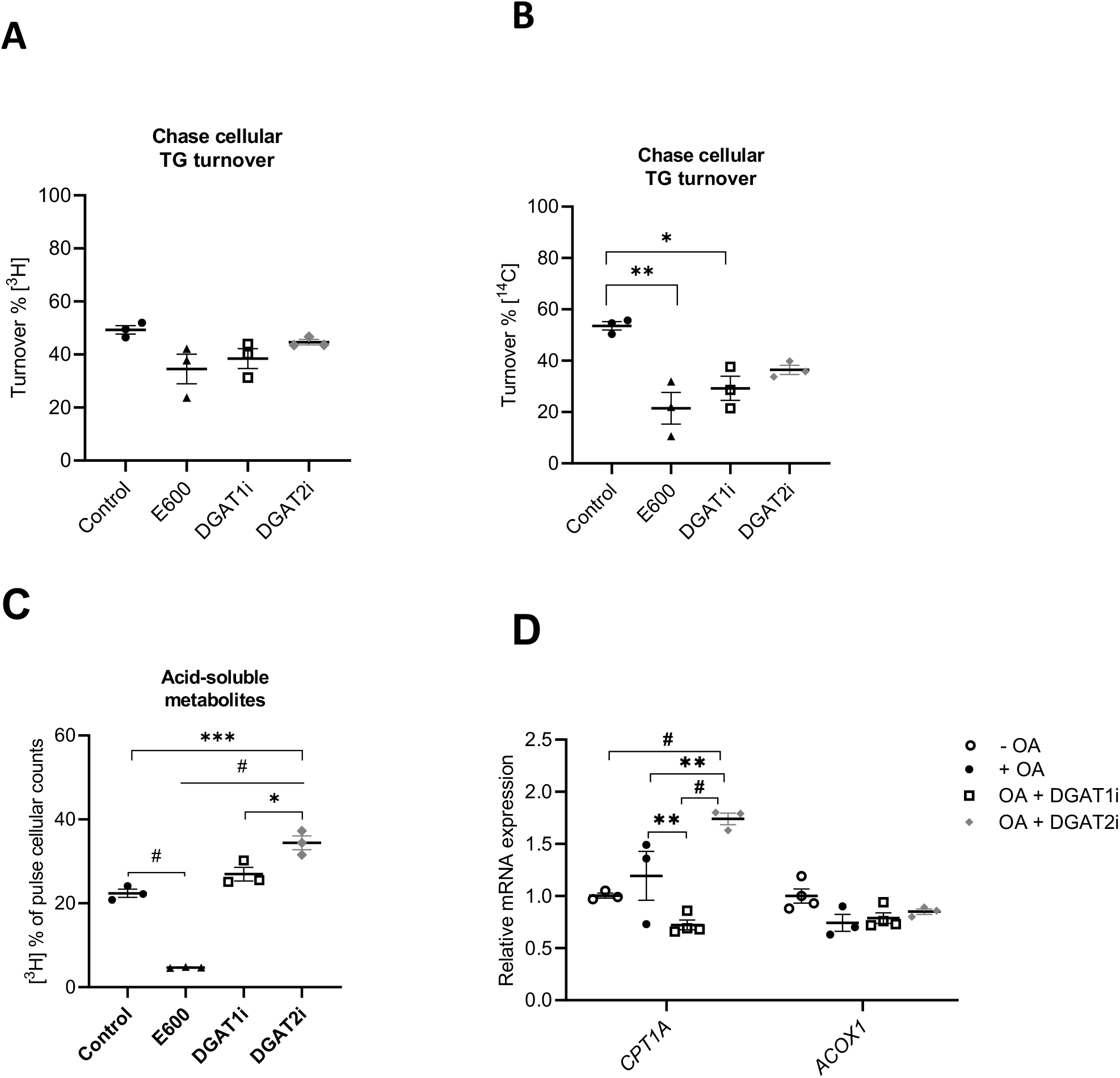
Turnover of DGAT1 and DGAT2 made TG. [^3^H]OA and [^14^C]glycerol dual labeled pulse-chase experiment was performed to assess turnover of TG in Huh7 hepatocytes in presence or absence of DGAT and lipase inhibitors. Percentage of turnover of preformed [^3^H] (**A**) and [^14^C] (**B**) cellular TG during the chase phase. (**C**) Acid-soluble metabolites produced from [^3^H]OA labeled lipids during chase. (D) mRNA expression of *CPT1A* and *ACOX1*.

### ATGL preferentially associates with DGAT1 made smaller LDs

Because DGAT1 synthesized TG appear to undergo increased turnover compared to DGAT2 generated TG we next investigated whether or not the LD lipolytic machinery including ATGL and ABHD5/CGI-58 preferentially targets to LDs containing DGAT1-made TG. Overexpression of active ATGL in Huh7 cells resulted in a dramatic depletion of LDs (Figure S3A) and therefore to investigate ATGL targeting to LDs we generated a catalytically inactive ATGL construct ATGL(S47A)-EGFP. ATGL(S47A)-EGFP predominantly localized on the LDs’ surface in OA treated Huh7 cells (Figure 5A). A significant increase in ATGL(S47A)-EGFP co-localization with LDs was observed in DGAT2 inhibitor treated cells compared with DGAT1 inhibitor treated cells, suggesting preferential ATGL targeting to DGAT1-made LDs (Figure 5A). ATGL is activated by ABHD5/CGI58. Therefore, we further analyzed localization of the ATGL activator on LDs generated by DGAT1 or DGAT2. Nearly all ABHD5/CGI58 localized to LDs and did not show preference for DGAT1- or DGAT2-made LDs (Figure 5B). To interrogate the colocalization between ATGL and ABHD5/CGI58 on LDs, we co-transfected cells with either ABHD5/CGI58-mCherry and active ATGL-EGFP or ABHD5/CGI58-mCherry and inactive ATGL(S47A)-EGFP followed by incubation with OA or OA+DGAT inhibitors. Co-transfection of active ATGL and ABHD5/CGI58 resulted in depletion of LDs to similar levels found in cells transfected with active ATGL alone (Figure S3B). The degree of inactive ATGL and ABHD5/CGI58 colocalization on LDs was not affected by either DGAT1 or DGAT2 inhibition (Figure 5C). Together, these data suggest that ATGL preferentially localizes to DGAT1-made LDs, and the ATGL activator ABHD5/CGI58 does not show this preference.

**Figure 5:**
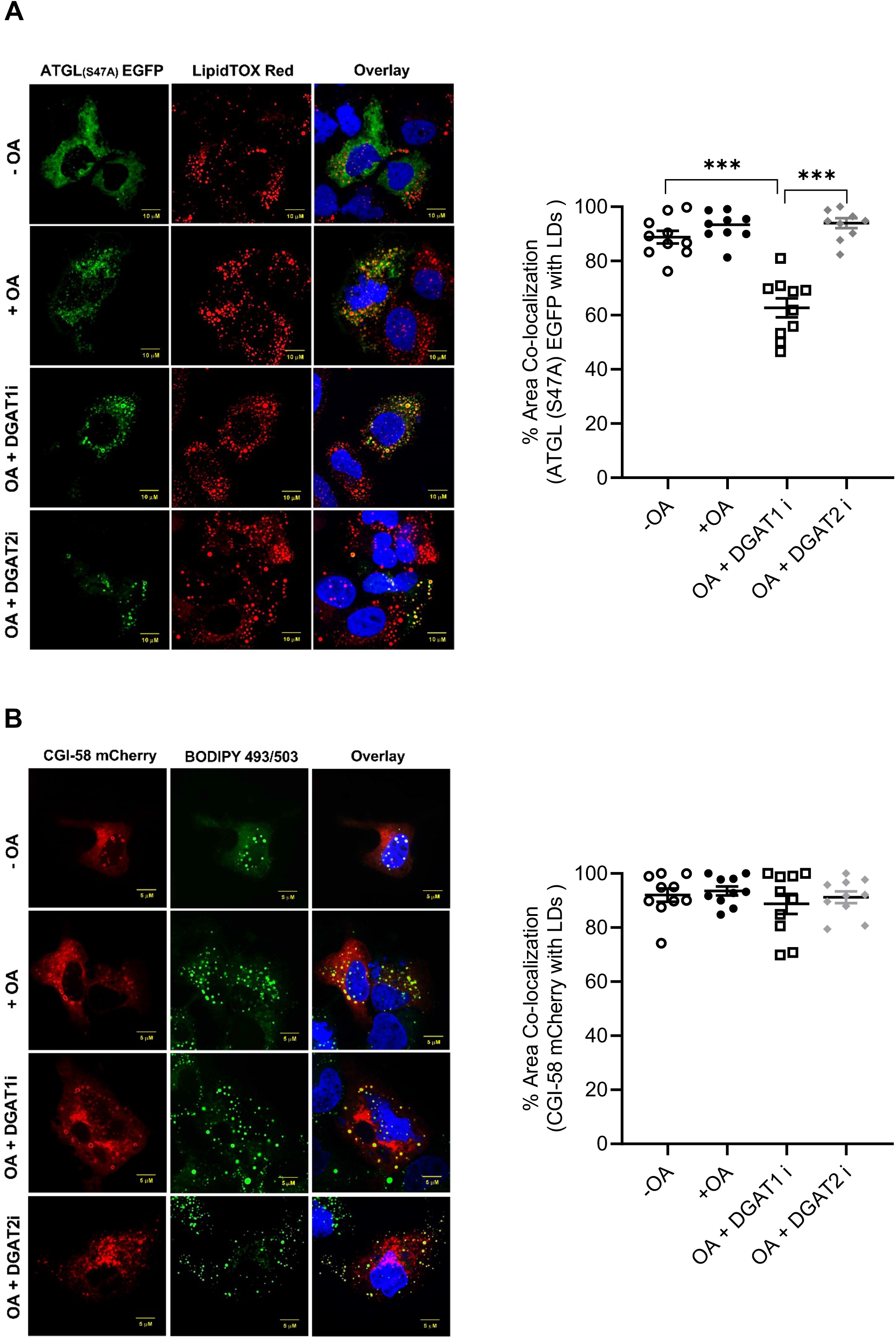
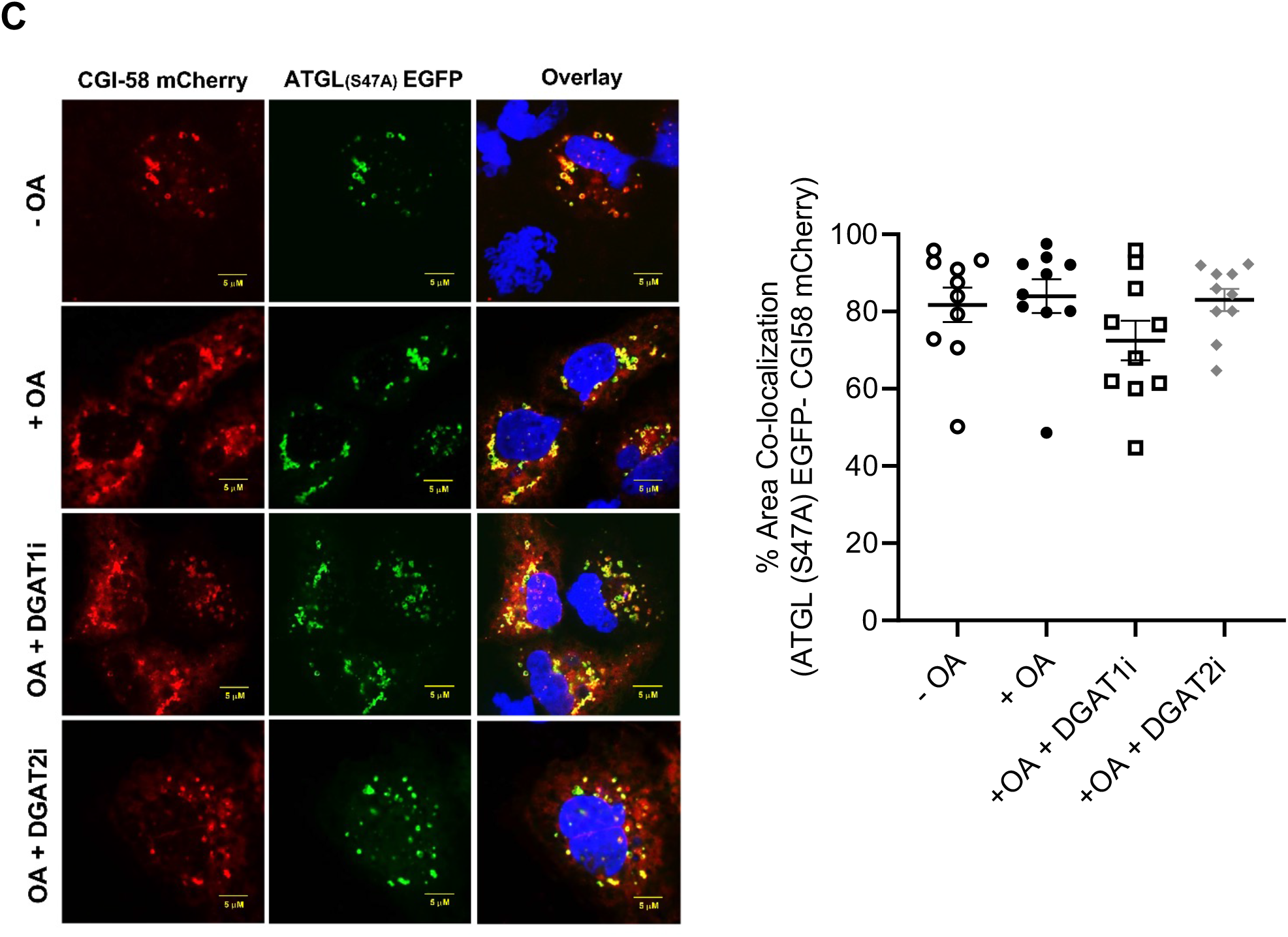
Effect of DGAT inhibitors on association of ATGL and ABHD5/CGI-58 with LDs. (**A**) Huh7 hepatocytes were transfected with ATGL(S47A)-EGFP and incubated with 0.4 mM OA/0.5% BSA for 4 h in the presence/absence of DGAT inhibitors. LDs were stained with LipidTox Red dye. **(B)** Huh7 hepatocytes were transfected with ABHD5/CGI58-mCherry and incubated with 0.4 mM OA/0.5% BSA for 4 h in the presence/absence of DGAT inhibitors. LDs were stained with BODIPY 493/503. (**C**) Huh7 hepatocytes were co-transfected with ATGL(S47A)-EGFP and ABHD5/CGI58-mCherry and incubated with 0.4 mM OA/0.5% BSA in the presence/absence of DGAT inhibitors for 4 h.

### DGAT inhibition does not affect PLIN2 targeting to LDs

PLIN2 and PLIN5 function as modulators of lipolysis in hepatocytes and coordinate recruitment of cytosolic lipases and co-factors (Granneman et al., 2009; Sztalryd and Brasaemle, 2017). Thus, we interrogated PLIN2 and PLIN5 targeting to LDs in the presence of DGAT1 or DGAT2 inhibitors. Inactivation of either DGAT1 or DGAT2 had no effect on targeting of transfected PLIN2-GFP to LDs (Figure 6A). On the other hand, co-localization of PLIN5-mCherry with LDs was reduced in cells treated with DGAT2 inhibitor compared to cells treated with DGAT1 inhibitor (Figure 6B). No differences were observed in the co-localization of inactive ATGL construct with PLIN5-mCherry in DGAT1 or DGAT2 inhibitor treated cells (Figure 6C). Together, these data suggest that DGAT inhibitors do not affect PLIN2 localization on LDs, whereas DGAT2i reduces the PLIN5 localization on LDs in Huh7 hepatocytes.

**Figure 6:**
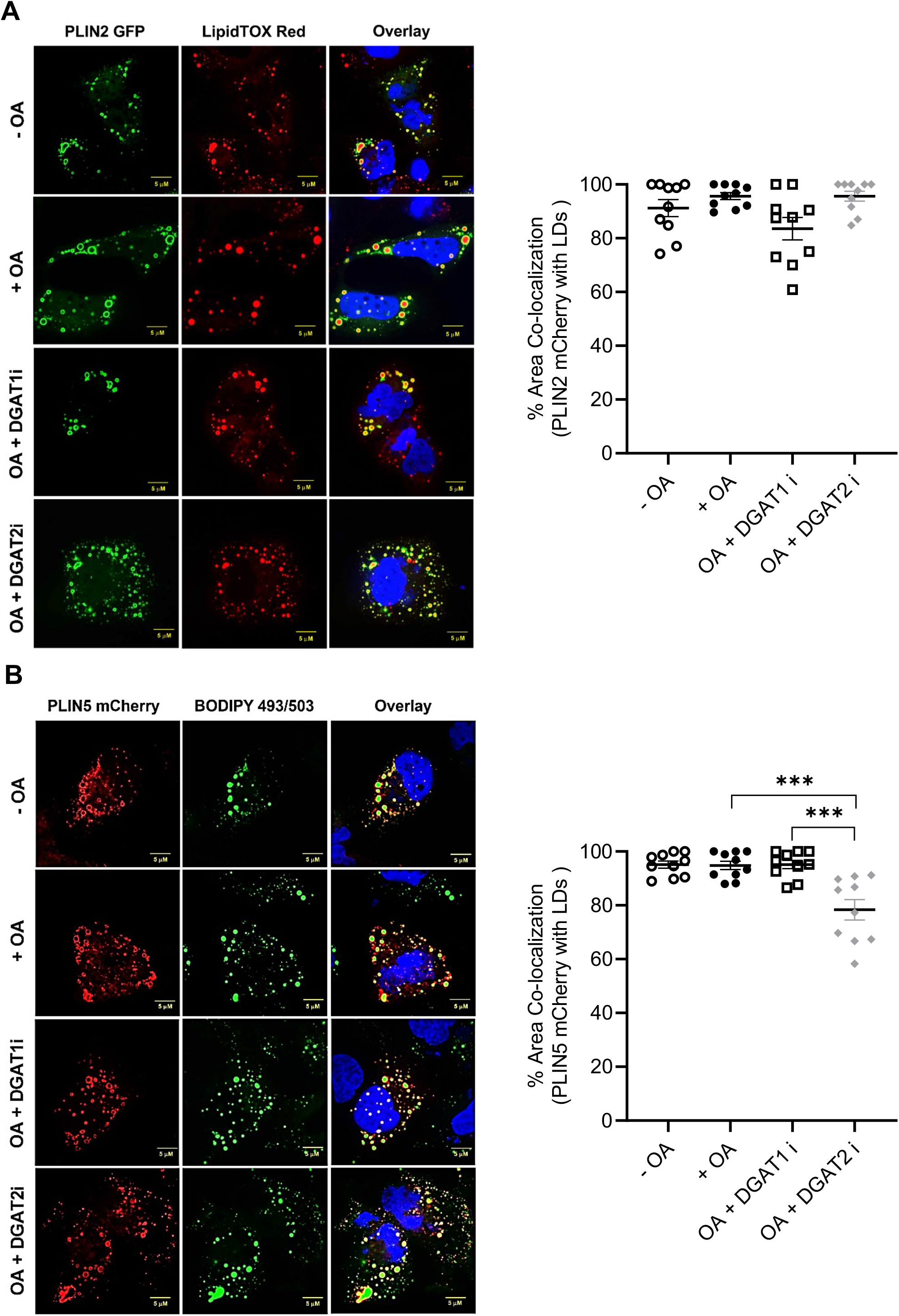
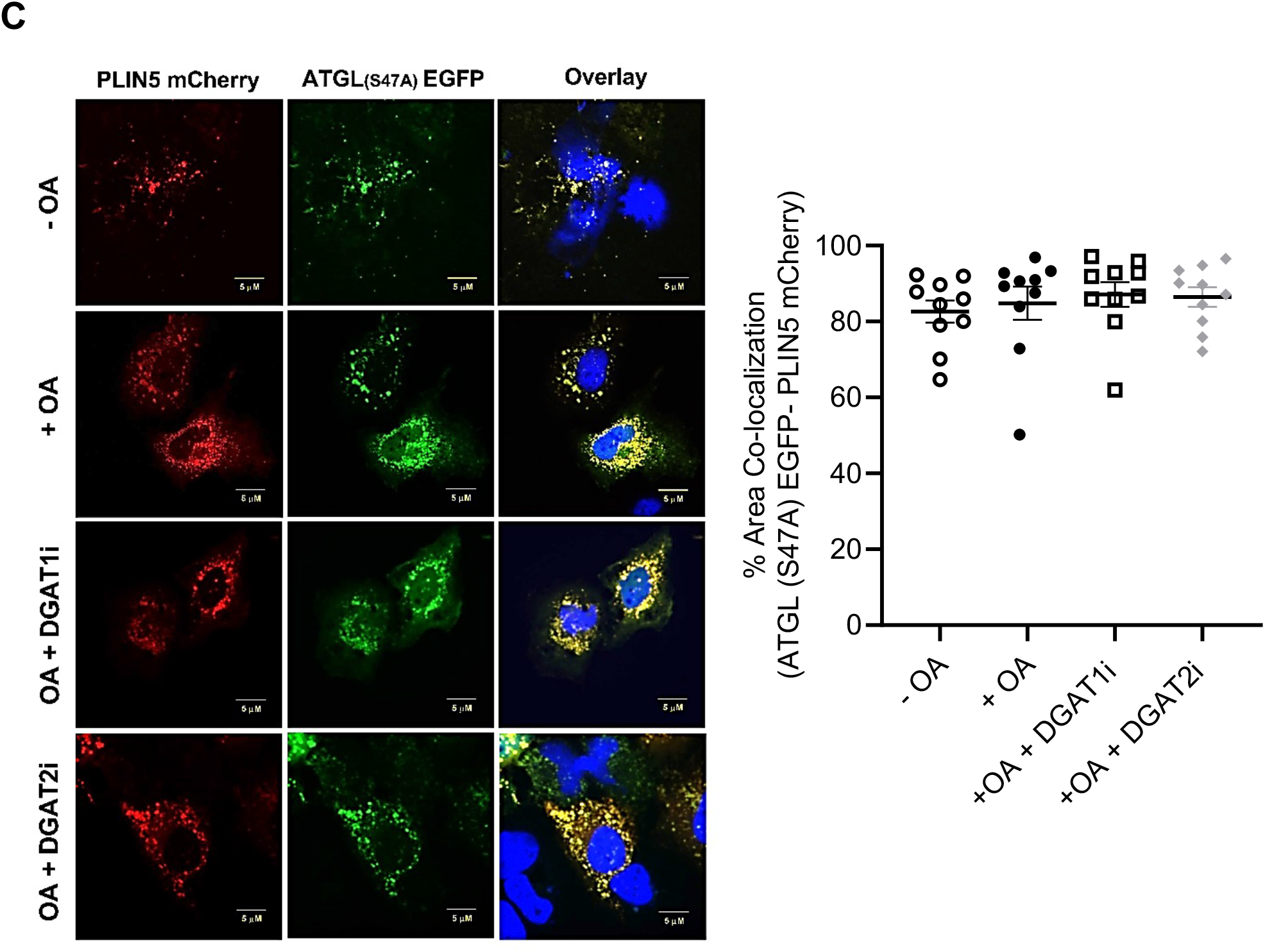
Effect of DGAT inhibition on targeting of PLIN2 and PLIN5 to LDs. **(A)** Huh7 hepatocytes were transfected with PLIN2-EGFP and incubated with 0.4 mM OA/0.5% BSA in the presence/absence of DGAT inhibitors for 4 h. LDs were stained with LipidTox Red dye. **(B)** Huh7 hepatocytes were transfected with PLIN5-mCherry and incubated with 0.4 mM OA/0.5% BSA in the presence/absence of DGAT inhibitors for 4 h. LDs stained with BODIPY 493/503. **(C)** Huh7 hepatocytes were co-transfected with ATGL(S47A)-EGFP and PLIN5-mCherry and incubated with 0.4 mM OA/0.5% BSA in the presence/absence of DGAT inhibitors for 4 h.

## Discussion

This study focused on delineating the role of DGAT1 and DGAT2 in TG accretion and utilization in the human hepatocyte model cell line Huh7. Our earlier studies have demonstrated that DGAT1 and DGAT2 can compensate for each other to synthesize triacylglycerol in primary mouse and human hepatocytes, and that DGAT1-made TG was deposited in small LDs, whereas DGAT2-made TG was stored in larger LDs (Li et al., 2015). However, the mechanisms giving rise to such different LD morphologies during incubation of cells with fatty acid are unclear. We observed similar LD morphological differences between DGAT1- and DGAT2-made LDs in Huh7 cells without any differences in TG mass. This suggests that either of the two DGATs can compensate for the loss of the other DGAT activity to protect the cell from lipotoxic accumulation of fatty acids. However, why does DGAT1 deposit TG in small sized LDs, while DGAT2 activity leads to formation of larger sized LDs? We tested the hypothesis that the smaller sized LDs generated by DGAT1 could be the result of increased turnover (lipolysis) rather than a specific mechanism that would prevent formation of larger LDs. Indeed, inhibition of lipolysis resulted in increased LD size of DGAT1-made LDs. On the other hand, inhibition of lipolysis in cells where TG synthesis was catalyzed by DGAT2 did not lead to increased LD size. These data suggest that DGAT1-made TG is more susceptible to lipolytic turnover compared to DGAT2-made TG. TG turnover studies confirmed increased oxidation of fatty acids originating from DGAT1-made TG. Increased turnover and fatty acid oxidation of TG synthesized by DGTA1 was accompanied by augmented abundance of *CPT1A* mRNA, further supporting the role of DGAT1-made TG as a source of fatty acid for energy production.

DGAT2 activity was found to be connected with de novo lipogenesis (Chitraju et al., 2017; Chitraju et al., 2019; Li et al., 2015; Villanueva et al., 2009; Wurie et al., 2012) and with regulation of a key transcription factor controlling lipogenesis SREBP1 (Choi et al., 2007). Inhibition of DGAT2 decreased *FASN* mRNA expression in Huh7 cells. Exogenous OA supplementation also reduced *SREBF1* and its target gene *SCD* mRNA expression.

The increased turnover of DGAT1 made TG compared to DGAT2 made TG suggests differential targeting of lipolytic machinery including ATGL and its activator ABHD5/CGI58 or inhibitory PLINs to the LDs. Previous studies have shown that overexpression of ATGL leads to smaller LDs, whereas knockdown or inhibition ATGL results in larger LDs in HeLa and AML12 cells (Schott et al., 2019; Smirnova et al., 2006). Specific ablation of ATGL activity in the mouse liver leads to TG storage in large LDs (Ong et al., 2011). In our studies overexpression of active ATGL (ATGL-EGFP) in Huh7 resulted in a complete absence of LDs, regardless whether TG was synthesized by DGAT1 or DGAT2. This suggests that either ATGL equally targeted to DGAT1 and DGAT2 generated LDs to hydrolyze TG, or the lipase hydrolyzed nascent LDs associated with the ER and mature LDs were not formed at all. To determine the ATGL and ABHD5/CGI58 preference for LDs containing TG made by DGAT1 or DGAT2 we used catalytically inactive ATGL(S47A)-EGFP to assess its targeting to LDs. ATGL(S47A)-EGFP localization percentage was higher on DGAT1 generated LDs compared to DGAT2-made LDs. This agrees with data demonstrating that fatty acids released through ATGL-mediated lipolysis are targeted for oxidation. Decreased localization of ATGL(S47A)-EGFP on DGAT2 formed LDs suggests that ATGL would not be involved in lipolysis to support the production of substrates for VLDL secretion. This data is consistent with studies from hepatocyte-specific deficient ATGL mice in which VLDL production is unaffected (Wu et al., 2011).

The lipase activity of ATGL is augmented through its association with the co-activator ABHD5/CGI58. ABHD5/CGI58 knockdown causes an abnormal accumulation of LDs in hepatocytes and preadipocytes (Yamaguchi et al., 2007). Ablation of ABHD5/CGI58 in mouse liver resulted in a substantial reduction of TG hydrolase activity and fatty acid oxidation, and facilitated TG and LD accumulations (Guo et al., 2013). As an extension of these studies, we sought to analyze the ABHD5/CGI58 localization on LDs generated by DGAT1 or DGAT2 activities. Given the increased presence of ATGL on DGAT1-made LDs and use of these LDs to support fatty acid oxidation, we also expected an augmented presence of ABHD5/CGI58 on these LDs. However, we found that ABHD5/CGI58 localized equally to both smaller and larger LDs, suggesting no preference of this lipolytic activator for either DGAT1- or DGAT2-made LDs.

PLIN1, PLIN2, and PLIN5 have been all documented to interact with ATGL and/or ABHD5/CGI58. (Granneman et al., 2011; Macpherson et al., 2013; Miyoshi et al., 2007; Yamaguchi et al., 2004). PLIN2 deficient mice present with reduced hepatic lipid accumulation and are resistant to diet-induced liver steatosis (Libby et al., 2016; McManaman et al., 2013; Najt et al., 2016). Importantly, the loss of lipase access barrier protection on LDs by PLIN2 resulted in increased lipolysis (Feng et al., 2017; Kimmel and Sztalryd, 2016). We found that PLIN2 expression (mRNA and protein) increased in Huh7 cells challenged with OA, which is consistent with a previous study in myoblasts where OA-mediated stimulation of PLIN2 expression was also observed (Xu et al., 2019). The abundance of PLIN2 plays an important role for the stability of expanded LDs (Thiam et al., 2013). Inhibition of DGAT1 or DGAT2 individually did not alter PLIN2 expression. This is consistent with other recent studies (Haaker et al., 2020; Xu et al., 2019). Combined DGAT1 and DGAT2 inhibition resulted in increased expression of *PLIN2* mRNA but did not change the PLIN2 protein levels. Similar results were obtained in *Dgat1*^-/-^/*Dgat2*^-/-^ mouse C_2_C_12_ myoblasts (Xu et al., 2019). OA was shown to increase both mRNA and protein expression of PLIN5 in AML12 and HepG2 cells (Li et al., 2012; Zhong et al., 2019). In contrast, PLIN5 expression (mRNA and protein) was unresponsive to OA supplementation in Huh7 cells and remained unchanged in the individual or combined DGAT inhibitor treatments. Both PLIN2 and PLIN5 targeted to DGAT1 or DGAT2 generated LDs without any obvious preference.

In conclusion, our data suggest that the smaller sized LDs observed in DGAT2 inhibited cells are a result of increased LD turnover through a lipase-dependent mechanism rather than by the inability of DGAT1 to support LD growth (Figure 7). Inhibition of DGAT1 or DGAT2 did not affect LD targeting of PLIN2 or ABHD5/CGI58, but inactivation of DGAT2 reduced PLIN5 targeting to LDs.

**Figure 7:**
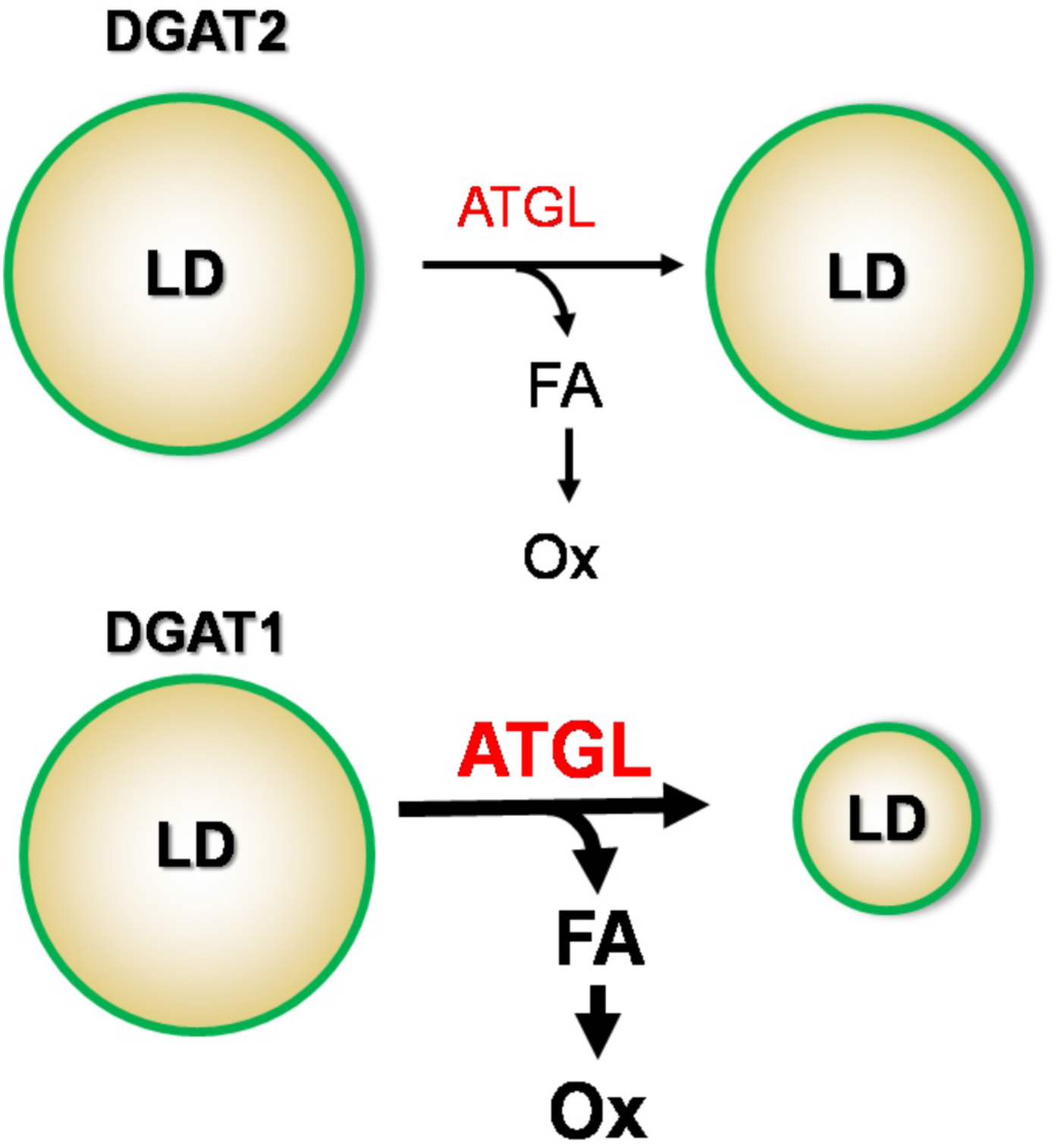
A working model of DGAT1 and DGAT2 contribution to LD storage and turnover in human hepatocytes. DGAT1 made LDs are preferentially targeted by the lipolytic machinery resulting in the reduction of LD size and increased provision of fatty acids for oxidation.

## Acknowledgments

This work was supported by the grants from Natural Sciences and Engineering Research Council RGPIN-2017-04734 and Canadian Institutes of Health Research PS 156314. We would like to thank Ms. Felicity Liu for her technical contribution to the study.

Confocal microscopy experiments were performed at the University of Alberta Faculty of Medicine & Dentistry Cell Imaging Centre, RRID:SCR_019200. Cell Imaging Centre receives financial support from the Faculty of Medicine & Dentistry, the Department of Medical Microbiology and Immunology, and Canada Foundation for Innovation (CFI) awards to contributing investigators. HPLC and GC lipid analyses were performed at the University of Alberta Faculty of Medicine & Dentistry Lipidomics Core, RRID:SCR_019176. Lipidomics Core receives financial support from the Faculty of Medicine & Dentistry, and Canada Foundation for Innovation (CFI) and Natural Sciences and Engineering Research Council of Canada (NSERC) awards to contributing investigators.

## References

Cases, S., Smith, S.J., Zheng, Y.W., Myers, H.M., Lear, S.R., Sande, E., Novak, S., Collins, C., Welch, C.B., Lusis, A.J., et al. (1998). Identification of a gene encoding an acyl CoA:diacylglycerol acyltransferase, a key enzyme in triacylglycerol synthesis. Proc Natl Acad Sci U S A 95, 13018–13023.

Chen, H.C., and Farese, R.V., Jr. (2005). Inhibition of triglyceride synthesis as a treatment strategy for obesity: lessons from DGAT1-deficient mice. Arterioscler Thromb Vasc Biol 25, 482–486.

Chen, H.C., Jensen, D.R., Myers, H.M., Eckel, R.H., and Farese, R.V., Jr. (2003). Obesity resistance and enhanced glucose metabolism in mice transplanted with white adipose tissue lacking acyl CoA:diacylglycerol acyltransferase 1. J Clin Invest 111, 1715–1722.

Chen, H.C., Stone, S.J., Zhou, P., Buhman, K.K., and Farese, R.V., Jr. (2002). Dissociation of obesity and impaired glucose disposal in mice overexpressing acyl coenzyme a:diacylglycerol acyltransferase 1 in white adipose tissue. Diabetes 51, 3189–3195.

Chitraju, C., Mejhert, N., Haas, J.T., Diaz-Ramirez, L.G., Grueter, C.A., Imbriglio, J.E., Pinto, S., Koliwad, S.K., Walther, T.C., and Farese, R.V., Jr. (2017). Triglyceride Synthesis by DGAT1 Protects Adipocytes from Lipid-Induced ER Stress during Lipolysis. Cell metabolism 26, 407–418.e403.

Chitraju, C., Walther, T.C., and Farese, R.V., Jr. (2019). The triglyceride synthesis enzymes DGAT1 and DGAT2 have distinct and overlapping functions in adipocytes. J Lipid Res 60, 1112–1120.

Choi, C.S., Savage, D.B., Kulkarni, A., Yu, X.X., Liu, Z.X., Morino, K., Kim, S., Distefano, A., Samuel, V.T., Neschen, S., et al. (2007). Suppression of diacylglycerol acyltransferase-2 (DGAT2), but not DGAT1, with antisense oligonucleotides reverses diet-induced hepatic steatosis and insulin resistance. J Biol Chem 282, 22678–22688.

Feng, Y.Z., Lund, J., Li, Y., Knabenes, I.K., Bakke, S.S., Kase, E.T., Lee, Y.K., Kimmel, A.R., Thoresen, G.H., Rustan, A.C., et al. (2017). Loss of perilipin 2 in cultured myotubes enhances lipolysis and redirects the metabolic energy balance from glucose oxidation towards fatty acid oxidation. J Lipid Res 58, 2147–2161.

Folch, J., Lees, M., and Sloane Stanley, G.H. (1957). A simple method for the isolation and purification of total lipides from animal tissues. J Biol Chem 226, 497–509.

Granneman, J.G., Moore, H.P., Krishnamoorthy, R., and Rathod, M. (2009). Perilipin controls lipolysis by regulating the interactions of AB-hydrolase containing 5 (Abhd5) and adipose triglyceride lipase (Atgl). J Biol Chem 284, 34538–34544.

Granneman, J.G., Moore, H.P., Mottillo, E.P., Zhu, Z., and Zhou, L. (2011). Interactions of perilipin-5 (Plin5) with adipose triglyceride lipase. J Biol Chem 286, 5126–5135.

Guo, F., Ma, Y., Kadegowda, A.K., Betters, J.L., Xie, P., Liu, G., Liu, X., Miao, H., Ou, J., Su, X., et al. (2013). Deficiency of liver Comparative Gene Identification-58 causes steatohepatitis and fibrosis in mice. J Lipid Res 54, 2109–2120.

Haaker, M.W., Kruitwagen, H.S., Vaandrager, A.B., Houweling, M., Penning, L.C., Molenaar, M.R., van Wolferen, M.E., Oosterhoff, L.A., Spee, B., and Helms, J.B. (2020). Identification of potential drugs for treatment of hepatic lipidosis in cats using an in vitro feline liver organoid system. Journal of veterinary internal medicine 34, 132–138.

Kimmel, A.R., and Sztalryd, C. (2016). The Perilipins: Major Cytosolic Lipid Droplet-Associated Proteins and Their Roles in Cellular Lipid Storage, Mobilization, and Systemic Homeostasis. Annual review of nutrition 36, 471–509.

Koliwad, S.K., Streeper, R.S., Monetti, M., Cornelissen, I., Chan, L., Terayama, K., Naylor, S., Rao, M., Hubbard, B., and Farese, R.V., Jr. (2010). DGAT1-dependent triacylglycerol storage by macrophages protects mice from diet-induced insulin resistance and inflammation. J Clin Invest 120, 756–767.

Li, C., Li, L., Lian, J., Watts, R., Nelson, R., Goodwin, B., and Lehner, R. (2015). Roles of Acyl-CoA:Diacylglycerol Acyltransferases 1 and 2 in Triacylglycerol Synthesis and Secretion in Primary Hepatocytes. Arteriosclerosis, thrombosis, and vascular biology 35, 1080–1091.

Li, H., Song, Y., Zhang, L.J., Gu, Y., Li, F.F., Pan, S.Y., Jiang, L.N., Liu, F., Ye, J., and Li, Q. (2012). LSDP5 enhances triglyceride storage in hepatocytes by influencing lipolysis and fatty acid β-oxidation of lipid droplets. PloS one 7, e36712.

Libby, A.E., Bales, E., Orlicky, D.J., and McManaman, J.L. (2016). Perilipin-2 Deletion Impairs Hepatic Lipid Accumulation by Interfering with Sterol Regulatory Element-binding Protein (SREBP) Activation and Altering the Hepatic Lipidome. J Biol Chem 291, 24231–24246.

Listenberger, L.L., Han, X., Lewis, S.E., Cases, S., Farese, R.V., Jr., Ory, D.S., and Schaffer, J.E. (2003). Triglyceride accumulation protects against fatty acid-induced lipotoxicity. Proc Natl Acad Sci U S A 100, 3077–3082.

Macpherson, R.E., Vandenboom, R., Roy, B.D., and Peters, S.J. (2013). Skeletal muscle PLIN3 and PLIN5 are serine phosphorylated at rest and following lipolysis during adrenergic or contractile stimulation. Physiological reports 1, e00084.

McFie, P.J., Banman, S.L., and Stone, S.J. (2018). Diacylglycerol acyltransferase-2 contains a c-terminal sequence that interacts with lipid droplets. Biochimica et biophysica acta. Molecular and cell biology of lipids 1863, 1068–1081.

McManaman, J.L., Bales, E.S., Orlicky, D.J., Jackman, M., MacLean, P.S., Cain, S., Crunk, A.E., Mansur, A., Graham, C.E., Bowman, T.A., et al. (2013). Perilipin-2-null mice are protected against diet-induced obesity, adipose inflammation, and fatty liver disease. J Lipid Res 54, 1346–1359.

Miyoshi, H., Perfield, J.W., 2nd, Souza, S.C., Shen, W.J., Zhang, H.H., Stancheva, Z.S., Kraemer, F.B., Obin, M.S., and Greenberg, A.S. (2007). Control of adipose triglyceride lipase action by serine 517 of perilipin A globally regulates protein kinase A-stimulated lipolysis in adipocytes. J Biol Chem 282, 996–1002.

Najt, C.P., Senthivinayagam, S., Aljazi, M.B., Fader, K.A., Olenic, S.D., Brock, J.R., Lydic, T.A., Jones, A.D., and Atshaves, B.P. (2016). Liver-specific loss of Perilipin 2 alleviates diet-induced hepatic steatosis, inflammation, and fibrosis. American journal of physiology. Gastrointestinal and liver physiology 310, G726–738.

Ong, K.T., Mashek, M.T., Bu, S.Y., Greenberg, A.S., and Mashek, D.G. (2011). Adipose triglyceride lipase is a major hepatic lipase that regulates triacylglycerol turnover and fatty acid signaling and partitioning. Hepatology 53, 116–126.

Paar, M., Jüngst, C., Steiner, N.A., Magnes, C., Sinner, F., Kolb, D., Lass, A., Zimmermann, R., Zumbusch, A., Kohlwein, S.D., et al. (2012). Remodeling of lipid droplets during lipolysis and growth in adipocytes. J Biol Chem 287, 11164–11173.

Schott, M.B., Weller, S.G., Schulze, R.J., Krueger, E.W., Drizyte-Miller, K., Casey, C.A., and McNiven, M.A. (2019). Lipid droplet size directs lipolysis and lipophagy catabolism in hepatocytes. The Journal of cell biology 218, 3320–3335.

Seebacher, F., Zeigerer, A., Kory, N., and Krahmer, N. (2020). Hepatic lipid droplet homeostasis and fatty liver disease. Seminars in cell & developmental biology.

Smirnova, E., Goldberg, E.B., Makarova, K.S., Lin, L., Brown, W.J., and Jackson, C.L. (2006). ATGL has a key role in lipid droplet/adiposome degradation in mammalian cells. EMBO Rep 7, 106–113.

Smith, S.J., Cases, S., Jensen, D.R., Chen, H.C., Sande, E., Tow, B., Sanan, D.A., Raber, J., Eckel, R.H., and Farese, R.V., Jr. (2000). Obesity resistance and multiple mechanisms of triglyceride synthesis in mice lacking Dgat. Nat Genet 25, 87–90.

Stone, S.J., Levin, M.C., Zhou, P., Han, J., Walther, T.C., and Farese, R.V., Jr. (2009). The endoplasmic reticulum enzyme DGAT2 is found in mitochondria-associated membranes and has a mitochondrial targeting signal that promotes its association with mitochondria. The Journal of biological chemistry 284, 5352–5361.

Stone, S.J., Myers, H.M., Watkins, S.M., Brown, B.E., Feingold, K.R., Elias, P.M., and Farese, R.V., Jr. (2004). Lipopenia and skin barrier abnormalities in DGAT2-deficient mice. J Biol Chem 279, 11767–11776.

Sztalryd, C., and Brasaemle, D.L. (2017). The perilipin family of lipid droplet proteins: Gatekeepers of intracellular lipolysis. Biochimica et biophysica acta. Molecular and cell biology of lipids 1862, 1221–1232.

Thiam, A.R., Farese Jr, R.V., and Walther, T.C. (2013). The biophysics and cell biology of lipid droplets. Nature Reviews Molecular Cell Biology 14, 775–786.

Villanueva, C.J., Monetti, M., Shih, M., Zhou, P., Watkins, S.M., Bhanot, S., and Farese, R.V., Jr. (2009). Specific role for acyl CoA:Diacylglycerol acyltransferase 1 (Dgat1) in hepatic steatosis due to exogenous fatty acids. Hepatology (Baltimore, Md.) 50, 434–442.

Walther, T.C., Chung, J., and Farese, R.V., Jr. (2017). Lipid Droplet Biogenesis. Annu Rev Cell Dev Biol 33, 491–510.

Walther, T.C., and Farese, R.V., Jr. (2012). Lipid droplets and cellular lipid metabolism. Annu Rev Biochem 81, 687–714.

Wang, H., Bell, M., Sreenivasan, U., Hu, H., Liu, J., Dalen, K., Londos, C., Yamaguchi, T., Rizzo, M.A., Coleman, R., et al. (2011). Unique regulation of adipose triglyceride lipase (ATGL) by perilipin 5, a lipid droplet-associated protein. J Biol Chem 286, 15707–15715.

Wilfling, F., Haas, J.T., Walther, T.C., and Farese, R.V., Jr. (2014). Lipid droplet biogenesis. Curr Opin Cell Biol 29, 39–45.

Wu, J.W., Wang, S.P., Alvarez, F., Casavant, S., Gauthier, N., Abed, L., Soni, K.G., Yang, G., and Mitchell, G.A. (2011). Deficiency of liver adipose triglyceride lipase in mice causes progressive hepatic steatosis. Hepatology 54, 122–132.

Wurie, H.R., Buckett, L., and Zammit, V.A. (2012). Diacylglycerol acyltransferase 2 acts upstream of diacylglycerol acyltransferase 1 and utilizes nascent diglycerides and de novo synthesized fatty acids in HepG2 cells. The FEBS journal 279, 3033–3047.

Xu, S., Zou, F., Diao, Z., Zhang, S., Deng, Y., Zhu, X., Cui, L., Yu, J., Zhang, Z., Bamigbade, A.T., et al. (2019). Perilipin 2 and lipid droplets provide reciprocal stabilization. Biophysics Reports 5, 145–160.

Yamaguchi, T., Omatsu, N., Matsushita, S., and Osumi, T. (2004). CGI-58 interacts with perilipin and is localized to lipid droplets. Possible involvement of CGI-58 mislocalization in Chanarin-Dorfman syndrome. J Biol Chem 279, 30490–30497.

Yamaguchi, T., Omatsu, N., Morimoto, E., Nakashima, H., Ueno, K., Tanaka, T., Satouchi, K., Hirose, F., and Osumi, T. (2007). CGI-58 facilitates lipolysis on lipid droplets but is not involved in the vesiculation of lipid droplets caused by hormonal stimulation. J Lipid Res 48, 1078–1089.

Yen, C.L., Stone, S.J., Koliwad, S., Harris, C., and Farese, R.V., Jr. (2008). Thematic review series: glycerolipids. DGAT enzymes and triacylglycerol biosynthesis. J Lipid Res 49, 2283–2301.

Zhong, W., Fan, B., Cong, H., Wang, T., and Gu, J. (2019). Oleic acid-induced perilipin 5 expression and lipid droplets formation are regulated by the PI3K/PPARα pathway in HepG2 cells. Applied physiology, nutrition, and metabolism = Physiologie appliquee, nutrition et metabolisme 44, 840–848.

Zimmermann, R., Strauss, J.G., Haemmerle, G., Schoiswohl, G., Birner-Gruenberger, R., Riederer, M., Lass, A., Neuberger, G., Eisenhaber, F., Hermetter, A., et al. (2004). Fat mobilization in adipose tissue is promoted by adipose triglyceride lipase. Science 306, 1383–1386.

