## Supplemental data for "Preferential lipolysis of DGAT1 over DGAT2 generated triacylglycerol in Huh7 hepatocytes"

**Supplemental Table S1. Primers used for quantitative PCR analysis**

| S.No | Gene | Primers |
|---|---|---|
| 1 | *ABHD5* | F: 5'- TTTAAGTCTAGTGCAGCGTTT – 3’ |
| | | R: 5’- GCATTTTACCAATTCGCTGGA – 3’ |
| 2 | *ACOX1* | F: 5'- TTTGAGGGAGAAAACACTGTCA – 3’ |
| | | R: 5’- CCTGAGTGCACCTGATCATAAC – 3’ |
| 3 | *BCSL2* | F: 5'- ATCTCCACTTCTTCGCGTTC – 3’ |
| | | R: 5’- CCAAATAGCAGGAGGCTAGAGA – 3’ |
| 4 | *CPT1A* | F: 5'- CCTCCGTAGCTGACTCGGTA – 3’ |
| | | R: 5’- CGGAGTGACCGTGAACTGA – 3’ |
| 5 | *PPIA* | F: 5'- TCCAAAGACAGCAGAAAACTTTCG – 3’ |
| | | R: 5’- TCTTCTTGCTGGTCTTGCCATTCC – 3’ |
| 6 | *DGAT1* | F: 5'- GTCCCAGGACTGCACCAG – 3’ |
| | | R: 5’- CTCCCCTCACACCACCAG – 3’ |
| 7 | *DGAT2* | F: 5'- TCGAGACTACTTTCCCATCCA – 3’ |
| | | R: 5’- GGTGGTATCCAAAGATATAGTTCCTG – 3’ |
| 8 | *FASN* | F: 5'- CAAGTGCACGGTGTTCCAT – 3’ |
| | | R: 5’- CTTCCCTTGGGCCATGTAG – 3’ |
| 9 | *FITM2* | F: 5'- AACAAGCGCAACGTCCTC – 3’ |
| | | R: 5’- TGGTAGTTGGTGAGGGCAAT– 3’ |
| 10 | *PLIN2* | F: 5'- GCTGCAGTCCGTCGATTT – 3’ |
| | | R: 5’- CACCCGAGTCACCACACTC – 3’ |
| 11 | *PLIN5* | F: 5'- CTTGAACACATGGACAAAGAGG – 3’ |
| | | R: 5’- TCAAGTTGGCCTGAATAGGG – 3’ |
| 12 | *SCD* | F: 5'- CCTAGAAGCTGAGAAACTGGTGA – 3’ |
| | | R: 5’- ACATCATCAGCAAGCCAGGT – 3’ |
| 13 | *SREBF1* | F: 5'- CGGAGCCATGGATTGCACTT – 3’ |
| | | R: 5’- TCAAATAGGCCAGGGAAGTCAC – 3’ |

#### Supplemental Data

##### Figure S1: *mRNA expression of genes encoding LD maturation factors*

Huh7 hepatocytes were incubated with/without 0.4 mM OA/0.5% BSA in the presence/absence of DGAT inhibitors for 4 h. Relative mRNA expression of *BSCL2* and *FITM2* encoding LD maturation factors SEIPIN and FIT2 (Fat storage-inducing transmembrane protein 2). Data are presented as the ratio of expression of a given gene to *PPIA* (cyclophilin).

##### Figure S2: *ATGL, CGI-58 and PLIN2 protein expression*

Huh7 hepatocytes were preincubated for 1 h with serum free DMEM incubated in the presence/absence of DGAT inhibitors followed by 4 h treatment with 0.4 mM OA/0.5% BSA and DGAT inhibitors in the presence/absence of lipase inhibitor E600. Immunoblot analysis of LD proteins, PLIN2, ATGL, ABHD5/CGI58, and GAPDH (loading control) were performed in total cell lysates.

##### Figure S*3: LD localization of ATGL-EGFP and co-localization of ATGL-EGFP and ABHD5/CGI58-mCherry*

(**A**) Huh7 hepatocytes were transfected with ATGL-EGFP and incubated with 0.4 mM OA/0.5% BSA for 4 h.

(**B**) Huh7 hepatocytes were co-transfected with ATGL-EGFP and ABHD5/CGI58-mCherry and incubated with 0.4mM OA/0.5% BSA for 4 h.

##### Figure S4: *Co-localization of ATGL and PLIN5*

(**A**) Huh7 hepatocytes were co-transfected with ATGL-EGFP and PLIN5-mCherry and incubated with 0.4mM OA/0.5% BSA for 4 h.

### Supplementary Figures

#### Figure S1

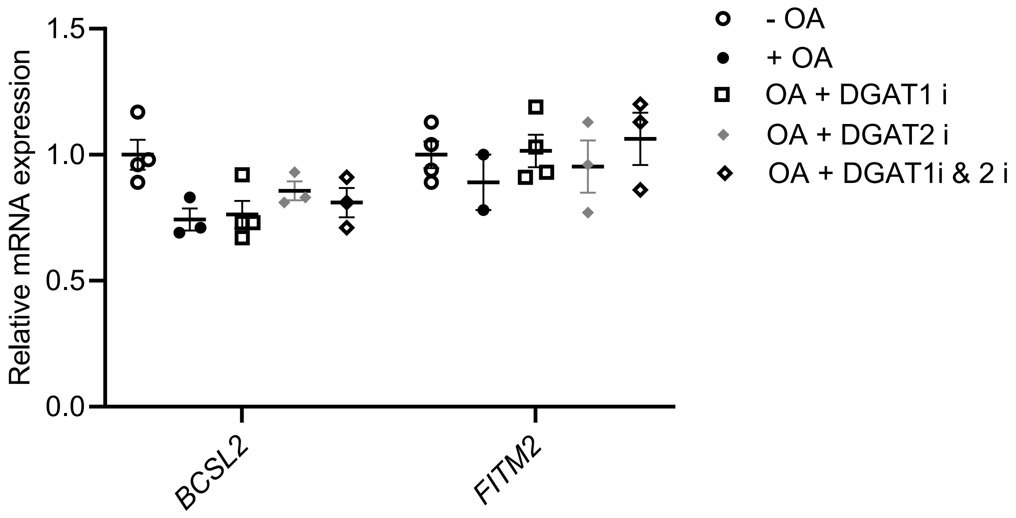

Figure S2

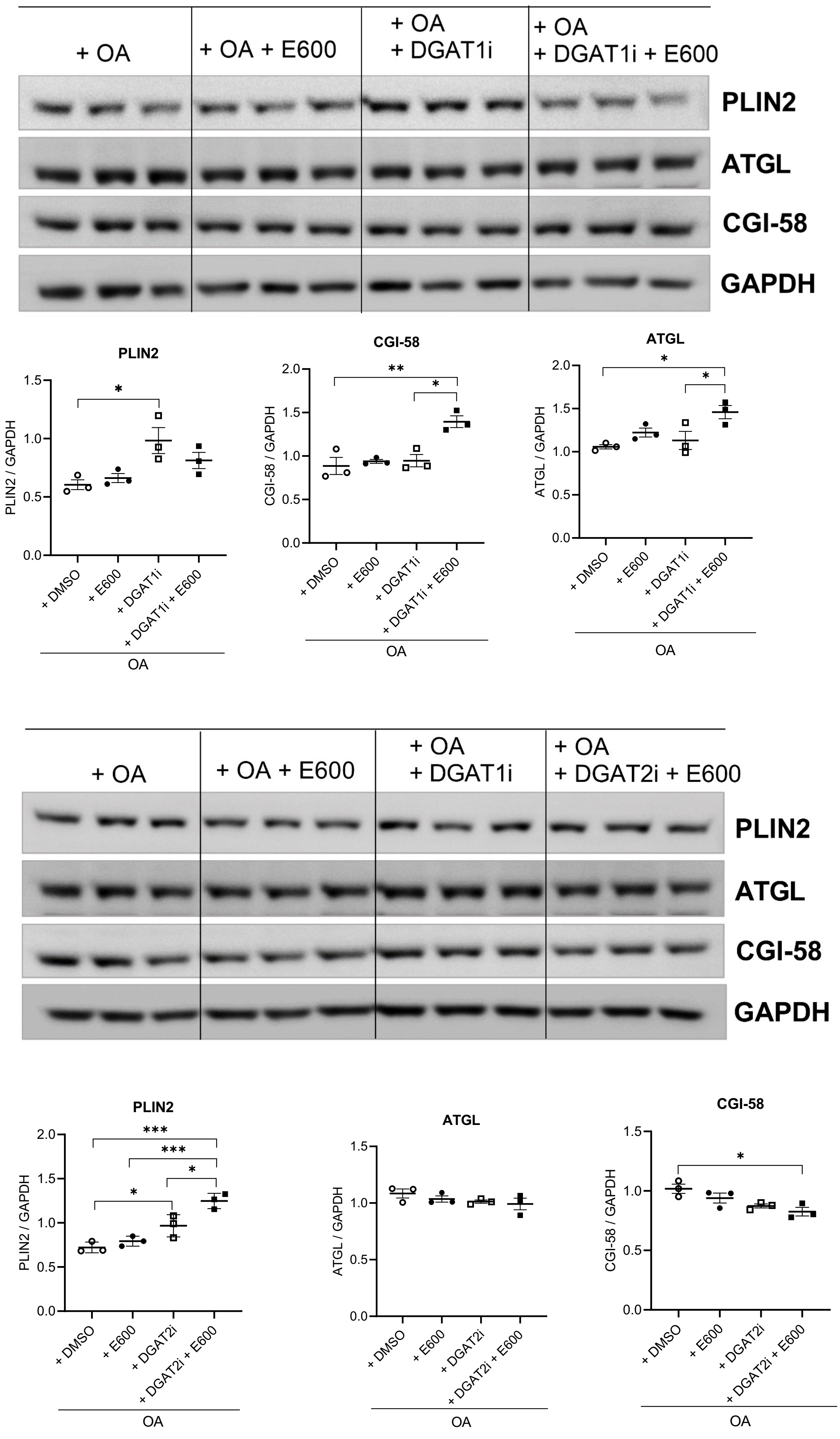
+ OA
+ OA + E600
+ OA
+ DGAT1i
+ OA
+ DGAT1i + E600
PLIN2
ATGL
CGI-58
GAPDH
PLIN2
PLIN2 / GAPDH
*
+ DMSO
+ E600
+ DGAT1i
+ DGAT1i + E600
OA
CGI-58
CGI-58 / GAPDH
**
*
ATGL
ATGL / GAPDH
+ OA
+ OA + E600
+ OA
+ DGAT1i
+ OA
+ DGAT2i + E600
PLIN2
ATGL
CGI-58
GAPDH
PLIN2
PLIN2 / GAPDH
***
+ DGAT2i
+ DGAT2i + E600
ATGL / GAPDH
CGI-58 / GAPDH

Figure S3

A

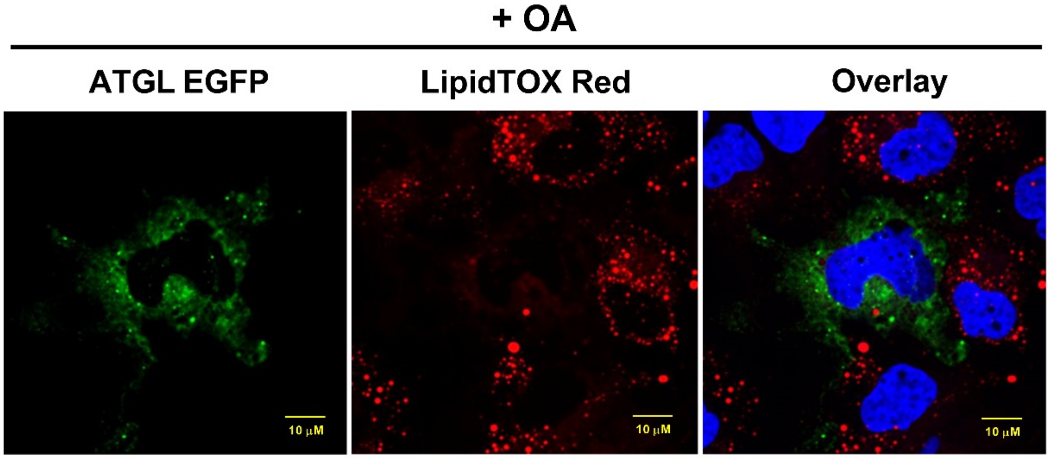
+ OA
ATGL EGFP
LipidTOX Red
Overlay
10 μM
10 μM
10 μM

B

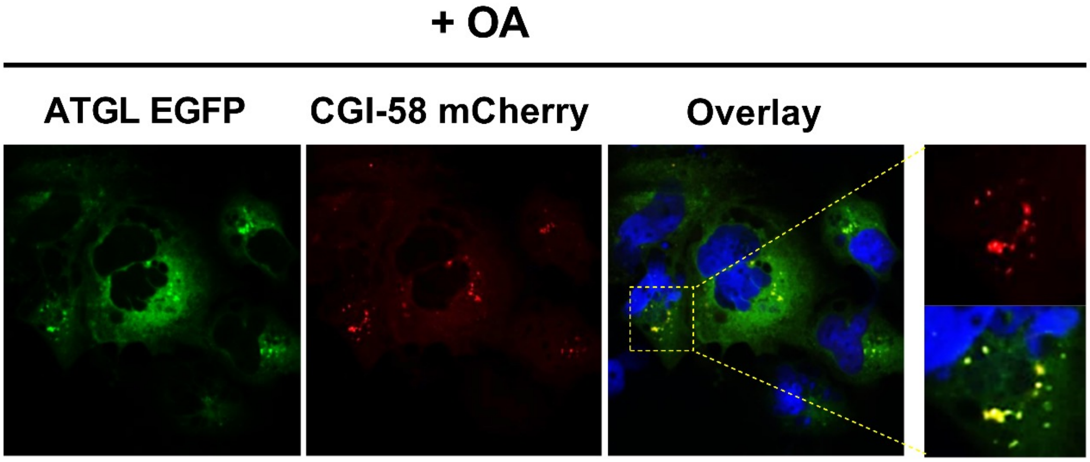
+ OA
ATGL EGFP
CGI-58 mCherry
Overlay

S4

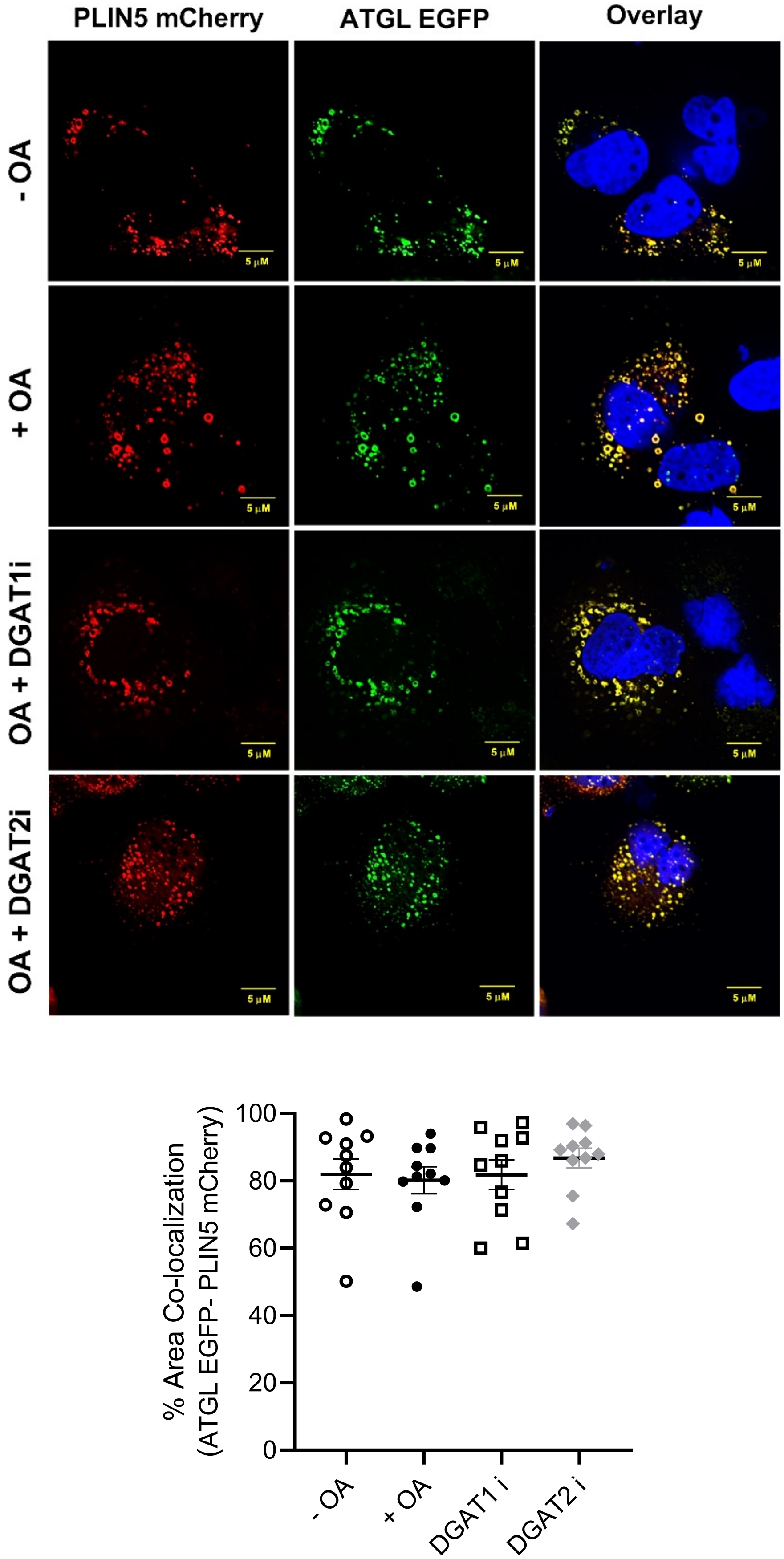

PLIN5 mCherry
ATGL EGFP
Overlay
- OA
+ OA
OA + DGAT1i
OA + DGAT2i
5 μM
% Area Co-localization
(ATGL EGFP- PLIN5 mCherry)
100
80
60
40
20
0
- OA
+ OA
DGAT1 i
DGAT2 i
